# Comprehensive analysis of mobile genetic elements in the gut microbiome reveals phylum-level niche-adaptive gene pools

**DOI:** 10.1101/214213

**Authors:** Xiaofang Jiang, Andrew Brantley Hall, Ramnik J. Xavier, Eric Alm

## Abstract

Mobile genetic elements (MGEs) drive extensive horizontal transfer in the gut microbiome. This transfer could benefit human health by conferring new metabolic capabilities to commensal microbes, or it could threaten human health by spreading antibiotic resistance genes to pathogens. Despite their biological importance and medical relevance, MGEs from the gut microbiome have not been systematically characterized. Here, we present a comprehensive analysis of chromosomal MGEs in the gut microbiome using a method called Split Read Insertion Detection (SRID) that enables the identification of the exact mobilizable unit of MGEs. Leveraging the SRID method, we curated a database of 5600 putative MGEs encompassing seven MGE classes called ImmeDB (Intestinal microbiome mobile element database) (<u>https://immedb.mit.edu/</u>). We observed that many MGEs carry genes that confer an adaptive advantage to the gut environment including gene families involved in antibiotic resistance, bile salt detoxification, mucus degradation, capsular polysaccharide biosynthesis, polysaccharide utilization, and sporulation. We find that antibiotic resistance genes are more likely to be spread by conjugation via integrative conjugative elements or integrative mobilizable elements than transduction via prophages. Additionally, we observed that horizontal transfer of MGEs is extensive within phyla but rare across phyla. Taken together, our findings support a phylum level niche-adaptive gene pools in the gut microbiome. ImmeDB will be a valuable resource for future fundamental and translational studies on the gut microbiome and MGE communities.

## Introduction

Horizontal gene transfer (HGT), the transfer of genes between organisms by means other than vertical transmission, allows for the rapid dissemination of genetic innovations between bacteria^1^. Ecology is an important factor shaping HGT, and the human gut in particular is a hotspot for HGT^2,3^. HGT impacts public health through its role in spreading antibiotic resistance genes^4,5^. The biological importance of HGT is exemplified by a porphyranase identified in *Bacteroides plebius* that digests seaweed, which was horizontally transferred from marine bacteria to human gut bacteria^6^. However, a major contributor to horizontal transfer - mobile genetic elements (MGEs) - have not been systematically characterized in the human gut microbiome.

Canonical classes of MGEs includes prophages^7^, group II introns^8^, and transposons^9^. It has become increasingly apparent that the acquisitions of a novel element class, genomic islands correspond to HGT events that differentiate commensal and pathogenic strains^10^. Genomic islands are non-canonical classes of MGEs that can transfer by conjugation or genomic regions derived from such MGEs. Integrative conjugative elements (ICEs) are a type of genomic island that can integrate into and excise from genomes using integrase, circularize using relaxase, replicate, and then transfer via conjugation^11,12^. Integrative mobilizable elements (IMEs) encode an integrase and relaxase for circularization like ICEs, but they have to hijack the conjugative machinery of co-resident ICEs or conjugative plasmids^13^.

Conventionally, HGTs are computationally identified by searching for the inconsistencies in the evolutionary history of gene and species^14^. However, this method overlooks the fact the horizontal transfer of multiple genes from the same locus might be the result of a single HGT event. Rather than individual genes, it is critical to identify the mobilizable units, in other words, the entire sequence of MGEs. Determining the mobilizable unit of MGEs is crucial to identify the mechanism of transfer, the preference of insertion sites, and cargo genes as well as to track the frequency of horizontal transfer events. In addition, information on MGEs are also valuable in the context of metagenomic analysis, as MGEs confound many metagenomics workflows such taxonomic profiling, strain-level variation detection, and pangenome analysis.

The repetitive and mobile nature of MGEs confounds many types of studies in microbiome communities, such as taxonomic profiling, strain-level variation detection, and pan-genome analyses. However, unlike research in eukaryotes, where multiple repeats databases exist for masking and annotation of repetitive DNA^15^, only a limited number of databases dedicated to the collection of MGE in prokaryotes^16–19^. Yet, these database are either limited one specific class of MGE or obsolete and not applicable for microbiome research. With the growing deluge of microbiome metagenomic sequencing data, a comprehensive MGE database of the gut microbiome is becoming increasingly critical.

In this study, we sought to characterize MGEs from the gut microbiome to understand how horizontal gene transfer by MGEs shapes the evolution of bacteria in the gut microbiome. First, we developed a method to identify the exact mobilizable unit of active MGEs using whole metagenome sequencing data together with references genomes. The algorithm implemented in SRID is similar to that of Daisy^20^, the first mapping-based HGT detection tool to our knowledge. Unlike Daisy, SRID was designed for use in a metagenomic context and doesn’t require pre-existing knowledge of both acceptor and donor genomes. We systematically identified MGEs with SRID and curated a database named ImmeDB (Intestinal microbiome mobile element database) dedicated to the collection, classification and annotation of these elements. The database is organized into seven MGE classes. Each MGE entry provides a visualization of annotations and downloadable genomic sequence and annotation. We detected many MGEs carrying cargo genes that confer an adaptive advantage to the gut environment. We also found that conjugation via integrative conjugative elements/ integrative mobilizable elements is more important than transduction via prophage for the spread of antibiotic resistance genes. This study provides insights into how the interplay of MGEs, bacteria, and the human host in the gut ecosystem lead to community-wide adaptations to the gut environment. The curated database of MGEs we have assembled here can be used by metagenomic workflows to improve future microbiome studies.

## Results

### Prevalence of MGEs in species of the gut microbiome

We systematically identified active MGEs from species of the human gut microbiome using mapping information from metagenomic reads from the Human Microbiome Project (HMP)^21^. MGEs are actively inserted and deleted from genomes, causing differences between strains of bacteria. We found cases where the reference genome of a bacterial strain differed from strains in the individual samples from the HMP. To find the sequences responsible for these differences, we mapped HMP metagenomic reads to available gut-associated bacterial reference genomes and identified genomic regions flanked by split reads and discordantly-aligned paired-end reads (Figure 1A). These regions potentially are recent insertions of active MGEs. The MGEs identified with the SRID method are limited to chromosomal MGEs. Thus, plasmids and extrachromosomal prophages were not characterized in this study. By searching for MGE-specific gene signatures, we verified and classified these MGEs (See Figure 1B and Methods).

**Figure 1.**
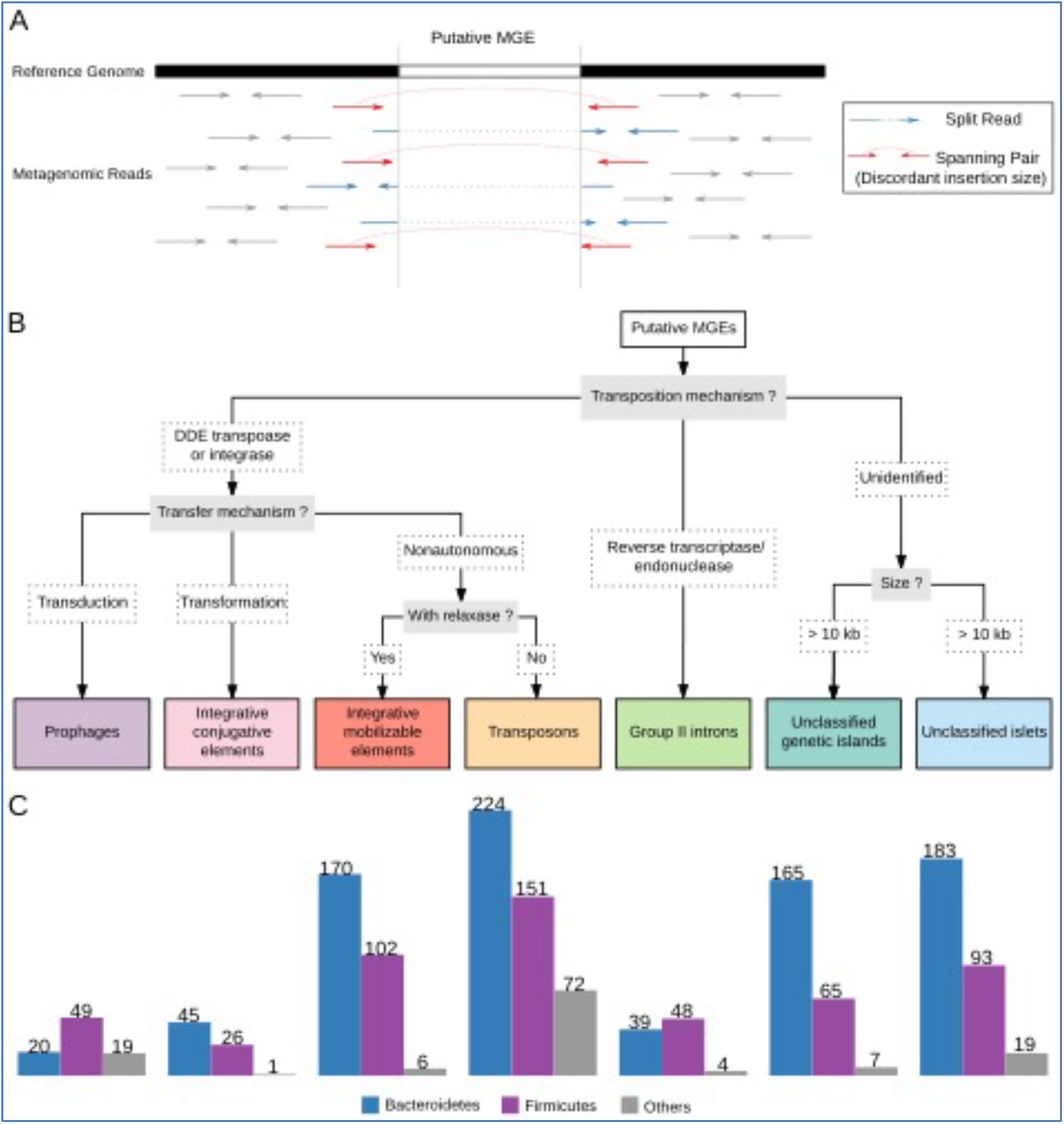
Identification and classification of gut microbiome MGEs. (A) The method used to identify putative MGEs using split reads and discordantly-mapped paired-end reads. Split reads are colored blue, and discordantly-mapped paired-end reads are colored red. (B) The method used to classify MGEs based on gene signatures. (C) The number of MGE clusters identified stratified by phyla and MGE classification.

We identified 5600 putative MGEs from gut microbiome representatives of 84 strains of Actinobacteria (10 species), 280 strains of Bacteroidetes (97 species), 158 strains of Firmicutes (118 species), 14 strains of Proteobacteria (12 species), and five strains of Verrucomicrobia (4 species) (Supplementary Table 2; Supplementary Data 1). Then, we classified the identified MGEs based on their transfer and transposition mechanisms into seven classes: ICEs, prophages, IMEs, group II introns, transposons, unclassified islets, and unclassified genomic islands (Figure 1C). Most of the MGEs identified (5145/5600) were from the phyla Bacteroidetes and Firmicutes because these two phyla tend to dominate the gut microbiome of healthy adults^21^. In general, smaller elements, such as transposons, had higher copy numbers per genome while larger elements, such as ICEs, prophages, and unclassified genomic islands, had a maximum of two copies per genome (Supplementary Figure 1).

Different strains of the same species often share identical or nearly-identical MGEs. To eliminate this redundancy, we collapsed MGEs into clusters based on overall nucleotide identity (Figure 1C). Phylum-level differences in the diversity of MGEs were revealed. For example, Bacteroidetes had more diversity of ICEs than Firmicutes (45 vs. 26 respectively), while Firmicutes had more diversity of prophages than Bacteroidetes (49 vs. 20 respectively).

### Diversity of MGE modules in gut microbiota

Although it has been known that ecology is important in shaping MGEs in the gut microbiome, this study is the first to systematically characterize the mechanisms of transposition and transfer for MGEs of the gut microbiome^2^. We annotated the genes in MGEs involved in their transposition and transfer, and then classified the elements into groups based on these annotations (Supplementary Table 2; Supplementary Data 2).

There are four major protein families responsible for transposition of gut MGEs: serine integrases, tyrosine integrases, DDE transposases, and group II intron proteins conferring reverse transcriptase and endonuclease activity. Serine and tyrosine integrases are the most prevalent protein families responsible for transposition in ICEs, IMEs, and prophages. In the gut microbiome MGE clusters we identified, we found 315 MGEs with tyrosine integrases (54 from ICEs, 206 from IMEs and 55 from prophages) and 110 MGEs with serine integrases (18 from ICEs, 67 from IMEs and 25 from prophages). Interestingly, while tyrosine integrases are found in several phyla, serine integrases of ICEs and prophages were exclusively found in the phylum Firmicutes. In IMEs, most serine integrases were identified in Firmicutes, but 10 clusters of serine integrases were found in Bacteroidetes and Actinobacteria (9 and 1 respectively). No ICEs and IMEs with DDE transposase were identified in our study. Nine prophage clusters were found with DDE transposase from IS families: IS30, IS256, and IS110. Interestingly, all three IS families use copy-paste mechanisms generating a transient double-stranded circular DNA intermediate to facilitate transposition^22^. This suggests that transient double-stranded circular intermediates may be essential for the life cycle of many prophages. All transposons we identified utilized DDE-transposase. We identified 19 families of transposase. Most of the transposase clusters we identified are present in insertion sequences. Seven clusters (28 copies) of transposons are composite transposons flanked by two different insertion sequences families.

ICEs and IMEs encode relaxases (MOB) to initiate DNA mobilization and transfer. We used the CONJscan-T4SSscan server to classify relaxases identified in MGEs^23^. Seven types of relaxase were identified in ICEs and IMEs. In ICEs, MOB_T_ was identified only in Firmicutes, MOB_V_ was identified only in Bacteroidetes, and MOBP1 was identified in Firmicutes, Bacteroidetes, and Actinobacteria. IMEs have a more diverse reservoir of relaxases. Besides the three types of relaxase found in ICEs, we also identified IMEs with MOB_P3_, MOB_B_, MOB_F_, and MOB_Q_ type relaxases.

ICEs are capable of conjugation via mating pair formation systems. Six types of mating pair formation systems for conjugation have been described^23^. We found three types of mating pair formation system: typeB, typeFA, and typeFATA, in ICEs from the gut microbiome. Consistent with previous findings, type FA systems were identified in 7 ICE clusters from Firmicutes, type B systems were identified in 45 ICE clusters from Bacteroidetes, and type FATA systems were identified in 19 Firmicutes ICE clusters and one Actinobacteria ICE cluster^24^.

### MGEs carry niche-adaptive genes

Although fundamentally selfish, MGEs often carry genes other than those necessary for their transposition and transfer, sometimes referred to as cargo genes^25^. We found that smaller elements like transposons generally carry zero or only a few cargo genes. Genetic islands like ICEs and IMEs often carry numerous cargo genes (median cargo genes 44 and 12 respectively). One example is an ICE found in *Bacteroides sp. 2_1_56FAA* (NZ_GL945043.1:1512740-1656974) which carries 139 cargo genes. We performed functional annotation on the cargo genes, and enrichment analysis using gene ontology (GO), Pfam, and Resfam^26–28^ (Supplementary Table 4). Several classes of enriched genes are well-known to be associated with the maintenance of MGEs such as restriction-modification systems and toxin-antitoxin pairs (Supplementary Table 4). Many other gene families carried by MGEs may confer an adaptive advantage to colonize the gut.

### Antibiotic resistance genes

Many classes of antibiotics consumed orally are incompletely absorbed in the small intestine, and therefore proceed to the large intestine where they can kill the resident microbes^29^. Therefore, genes that confer antibiotic resistance can be adaptive to the gut environment. In total, we identified 781 antibiotic resistance genes encompassing 46 distinct classes carried by MGEs. Classes of MGEs varied in their carriage of antibiotic resistance genes. Of 8151 prophage cargo genes, only 13 were found to be antibiotic resistance genes. The carriage rate of antibiotic resistance genes normalized by total cargo genes in prophages is more than ten times lower than that identified in ICEs (330/16820) and IMEs (229/11053) (Supplementary Figure 1). This suggests that conjugation via ICE/IME may be more important than transduction in the spread of antibiotic resistance genes, consistent with previous findings^30,31^.

GO analysis revealed that cargo genes from the class “rRNA modification” (GO:0000154), which confers resistance to a wide range of antibiotics including tetracycline and erythromycin, are enriched in both Bacteroidetes and Firmicutes. Resfam enrichment analysis also supported this, as RF0135 (tetracycline resistance ribosomal protection protein), and RF0067 (Emr 23S ribosomal RNA methyltransferase) were enriched. Other enriched antibiotic resistance gene classes carried by MGEs confer resistance to chloramphenicol (RF0058), cephalosporins (RF0049) and aminoglycosides (RF0167).

One example of an MGE responsible for the transmission of antibiotic resistance is the ICE CTnDOT, the spread of which dramatically increased the prevalence of tetracycline-resistant Bacteroidetes species^32^. CTnDOT-like ICEs were clustered in ICE1. Elements in this cluster typically confer resistance to tetracycline via the tetQ antibiotic resistance gene (Figure 2A). In addition, ICE1 elements have multiple sites where antibiotic resistance genes can be inserted or substituted. We characterized 5 insertions of antibiotic resistance genes into ICE1 (Figure 2A). Insertion sites 1, 2, and 5 are between operons; therefore they do not interrupt the function of crucial genes. We observed one insertion and two substitutions of antibiotic resistance genes around the tetQ operon, suggesting that this site is likely a “hotspot” for insertions and substitutions of antibiotic resistance genes. Our analysis reveals the surprising extent to which MGEs in species of the gut microbiome contribute to the phenomenon of antibiotic resistance and that the insertion of antibiotic resistance genes into MGEs is an active and ongoing process.

**Figure 2.**
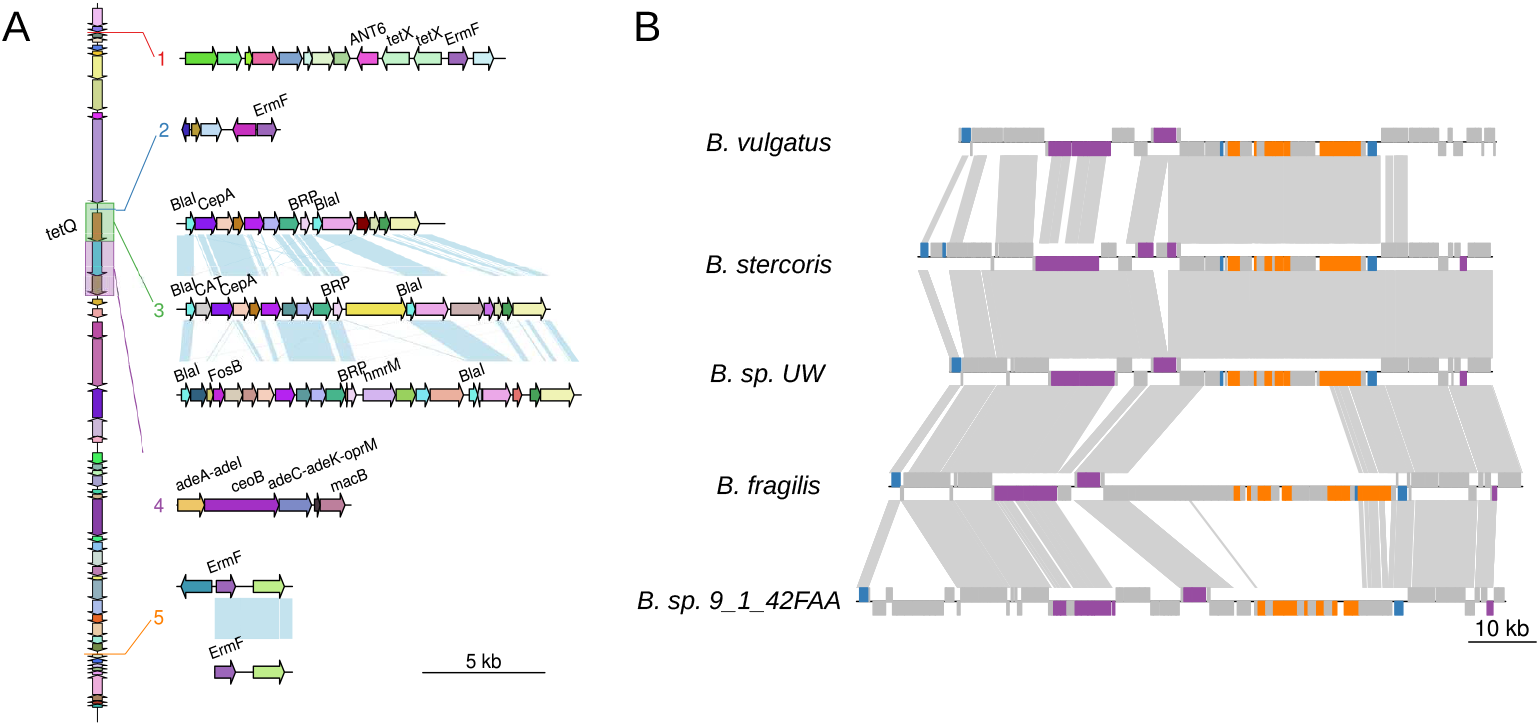
Examples of niche-adaptive genes. (A) CTn-DOT-like elements have acquired antibiotic resistance genes on multiple, independent occasions. Here, we show insertion sites of antibiotic resistant genes in CTnDOT-like elements. A CTnDOT-like ICE is shown on the left. Orthologs between elements are visualized using genoPlotR (light blue connections) and are the same color. Numbers in the top panel represent the insertion site of the numbered elements below. Antibiotic resistance genes are labeled. (B) ICEs are involved in the transfer of capsular polysaccharide biosynthesis loci between Bacteroidetes species. Here, we show examples of ICEs containing capsular polysaccharide biosynthesis loci. Orthologs between elements are plotted with GenoPlotR. Genes involved in capsular polysaccharide biosynthesis are colored orange, integrases are colored blue, and genes involved in conjugation are colored purple. Grey links indicate orthologs between elements.

### Bile salt hydrolase and bile transporters

Bile acids are found in high concentrations in the human intestines^33^ and can be toxic to bacteria^34^. Therefore, gut microbes have developed strategies to deal with bile acids by actively pumping bile acids out of the cell, or via deconjugation, which is hypothesized to diminish the toxicity of bile acids^33,34^. The high identity of archaeal and bacterial bile salt hydrolases strongly suggests the horizontal transfer of this gene^35^. A sodium bile acid symporter family (PF01758), which could help to pump bile acids out of the cell, was found to be enriched in the cargo genes of MGEs. Furthermore, 61 examples of bile salt hydrolases were identified as cargo genes of MGEs (Supplementary Table 3). Thus, MGEs carry genes that help microbes to overcome a specific challenge of colonizing the human gut.

### Glycoside hydrolases for mucus utilization

The colon is lined with a layer of mucus composed of the glycoprotein MUC2^36^. The glycans that decorate MUC2 have a core structure composed of galactose, N-acetylglucosamine, N-acetylgalactosamine, with terminal residues of fucose and sialic acid^37^. These specific glycans are a major energy source for members of the gut microbiota^38^. Therefore, it may benefit members of the gut microbiota to degrade these specific glycans^39^. We found cargo genes carried by MGEs from Bacteroidetes species were enriched for GO:0004308, an exo-sialidase involved in the degradation of mucosal glycans. In addition, we identified 60 glycoside hydrolases capable of degrading mucosal glycans carried by MGEs from the categories: sialidases (GH33), fucosidases (GH95), α-N-acetylgalactosaminidases (GH109), and β-galactosidases (GH20)^38,40^ (Supplementary Table 3). Thus, MGEs carry genes to unlock a key energy source available to gut microbes.

### Polysaccharide Utilization Loci

Gut Bacteroidetes can utilize a wide variety of polysaccharides via the products of polysaccharide utilization loci, which collectively make up large proportions of Bacteroidetes genomes^41^. Each polysaccharide utilization locus contains a copy of the gene SusC, a sugar transporter, and SusD, a glycan binding protein^42^. Due to the wide range of polysaccharides available to gut microbes, it is hypothesized that the possession of a large repertoire of polysaccharide utilization loci confers an adaptive advantage in Bacteroidetes^41^. We found 43 polysaccharide utilization loci containing both SusC and SusD carried by MGEs suggesting that the ability to degrade complex polysaccharides may be readily transferred between members of the gut microbiota (Supplementary Table 3).

### Capsular Polysaccharide Biosynthesis Loci

Many bacterial species produce capsules, an extracellular structure made up of polysaccharides^43^. However, gut Bacteroidetes species have a large repertoire of capsular polysaccharide biosynthesis loci (up to 8) compared to other bacterial species and even Bacteroidetes from other sites such as the mouth^44^. Furthermore, capsular polysaccharide biosynthesis loci have been reported to be the most polymorphic region of *Bacteroides* genomes^45,46^. Multiple capsular polysaccharide biosynthesis loci are necessary to competitively colonize the gut, and are therefore considered to be gut adaptive genes in gut Bacteroidetes^47^.

Capsular polysaccharide biosynthesis loci are large and complex; many contain upwards of 20 genes^43^. We found 21 complete or fragmented capsular polysaccharide biosynthesis loci containing at least 10 genes carried by MGEs (Supplementary Table 3). For example, almost identical copies of ICE9 containing a capsular polysaccharide biosynthesis locus were found in two species, *B. stercoris* and *B. sp. UW*. The same capsular polysaccharide biosynthesis locus was also found in *B. vulgatus*, but the ICE9 copy was slightly divergent. Two other copies of ICE9 likely containing an orthologous capsular polysaccharide biosynthesis locus were also found in *B. fragilis* and *B. sp. 9_1_42FAA* (Figure 2B). Additionally, many GO-terms related to capsular polysaccharide biosynthesis are enriched in Bacteroidetes MGEs including GO:0045226, GO:0034637, GO:0044264, and GO:0000271. A Pfam for glycosyltransferases involved in the biosynthesis of capsular polysaccharides (PF13579) was enriched in Bacteroidetes MGEs. The transfer of large segments of capsular polysaccharide biosynthesis loci by MGEs may help to explain the incredible diversity of capsular polysaccharide biosynthesis loci observed in the genomes of gut Bacteroidetes^48^.

### Sporulation

The gut is an anaerobic environment colonized by many classes of strictly anaerobic organisms^49,50^. However, to transmit between hosts, gut microbes must be exposed to oxygen. Recent work has shown that many more gut microbes form spores than previously thought, likely enabling transmission between hosts^51^. In Firmicutes, 14 genes involved in sporulation (GO:0030435) were found to be enriched in MGEs. In addition, PF08769 (Sporulation initiation factor Spo0A C terminal) and PF04026 (SpoVG) were also enriched in our Pfam analysis.

One example is GI153, a genetic island from *Faecalibacterium prausnitzii A2-165*, which contains a series of spore formation-related genes in an operon: SpoVAC, SpoVAD, spoVAEb, gpr (spore protease), and spoIIP. Another example is GI175, a genetic island derived from a degenerate prophage in *Roseburia intestinalis* L1-82. In one operon of GI175, there are three genes: SpoVAEb, SpoVAD, and one unknown gene with Cro/C1-type HTH DNA-binding domain. SpoVAC, SpoVAD, spoVAEb homologs were previously found to be carried by a Tn1546-like ICE and conferred heat resistance to spores in the model spore forming organism *Bacillus subtilis*^52^. Thus, MGEs may help to transfer genes involved in sporulation between gut microbiota which may prove adaptive for colonizing new hosts.

### Summary of cargo genes

Many additional gene families were found to be enriched in MGEs that could plausibly be niche adaptive including: histidine sensor kinases, and genes involved in vitamin B biosynthesis (Supplementary Table 4). Notably, MGEs from Firmicutes and Bacteroidetes have different types of genes enriched reflecting the differences in physiology between the phyla. Antibiotic resistance genes and genes involved in the detoxification of bile acids are enriched in MGEs from both phyla. Glycoside hydrolases for mucus utilization, and capsular polysaccharide biosynthesis loci are enriched only in MGEs from Bacteroidetes, while genes for sporulation are enriched in MGEs only from Firmicutes. Overall, the transfer of niche adaptive genes by MGEs likely has a large impact on the fitness of species of the gut microbiome.

### Host ranges and evolution of MGEs

Although MGEs readily transfer between species, there has not been a systematic analysis of the host range of MGEs in the gut microbiome. The host ranges of different classes of MGEs is variable, and even within a class, different elements have variable host ranges. Understanding the host range of gut MGEs is of particular importance because gut MGEs carry many cargo genes, and the host range of the MGE defines how widely these cargo genes can be distributed. For example, the gut microbiome is a reservoir of antibiotic resistance genes, and many antibiotic resistance genes are located within MGEs^53^. Therefore, it is important to understand the probability of the transfer of MGEs with antibiotic resistance genes from commensals to pathogens^53^.

First, we studied the host range of MGEs from the same cluster. MGEs in the same cluster that exist in at least two species generally represent recent horizontal transfer. Some MGE clusters are present in a wide range of species indicative of active horizontal transfer. One example is the ICE1 cluster, a representative of the CTnDOT-like ICEs, which is found in 32 species of Bacteroidetes from the genera: *Bacteroides, Parabacteroides, Allistipes*, and *Paraprevotella* (Supplementary Figure 1). The entirety of the 49kb element is found at more than 99 percent nucleotide identical to 10 *Bacteroides, Parabacteroides, and Allistipes* species, indicative of very recent horizontal transfer. This cluster also includes other CTnDOT-like elements with more variability such as CTnERL, which has an additional insertion of an IME conferring erythromycin resistance^54^. Another example is the Firmicutes ICE cluster ICE10, which is found in 10 species of the families *Lachnospiraceae* and *Ruminococcaceae*. This ICE10 cluster belongs to Tn916/Tn1549 family of ICEs, some members of which carry the medically-important VanB gene conferring resistance to vancomycin^55^. We found no examples of ICEs from the same cluster present in multiple phyla. Clusters of prophage, IMEs, group II introns, and transposons were also found in many species but were again limited to a single phylum. Our results support that although the recent horizontal transfer of MGEs is common within phyla, cross-phyla horizontal transfer is rare, as we did not observe any cross-phyla horizontal transfer events for elements of the same cluster.

Here we generated phylogenetic trees of tyrosine and serine integrases from ICEs and prophages identified to study the evolutionary history of the recombination module of MGEs. To contrast the phylogeny of the tyrosine and serine integrases with host species lineages we plotted tanglegrams (Figure 3 and Figure 4). The phylogeny of both the serine and tyrosine integrases is incongruent with the host species lineages which is indicative of extensive past horizontal transfer of ICEs and prophages between species of the gut microbiome.

**Figure 3.**
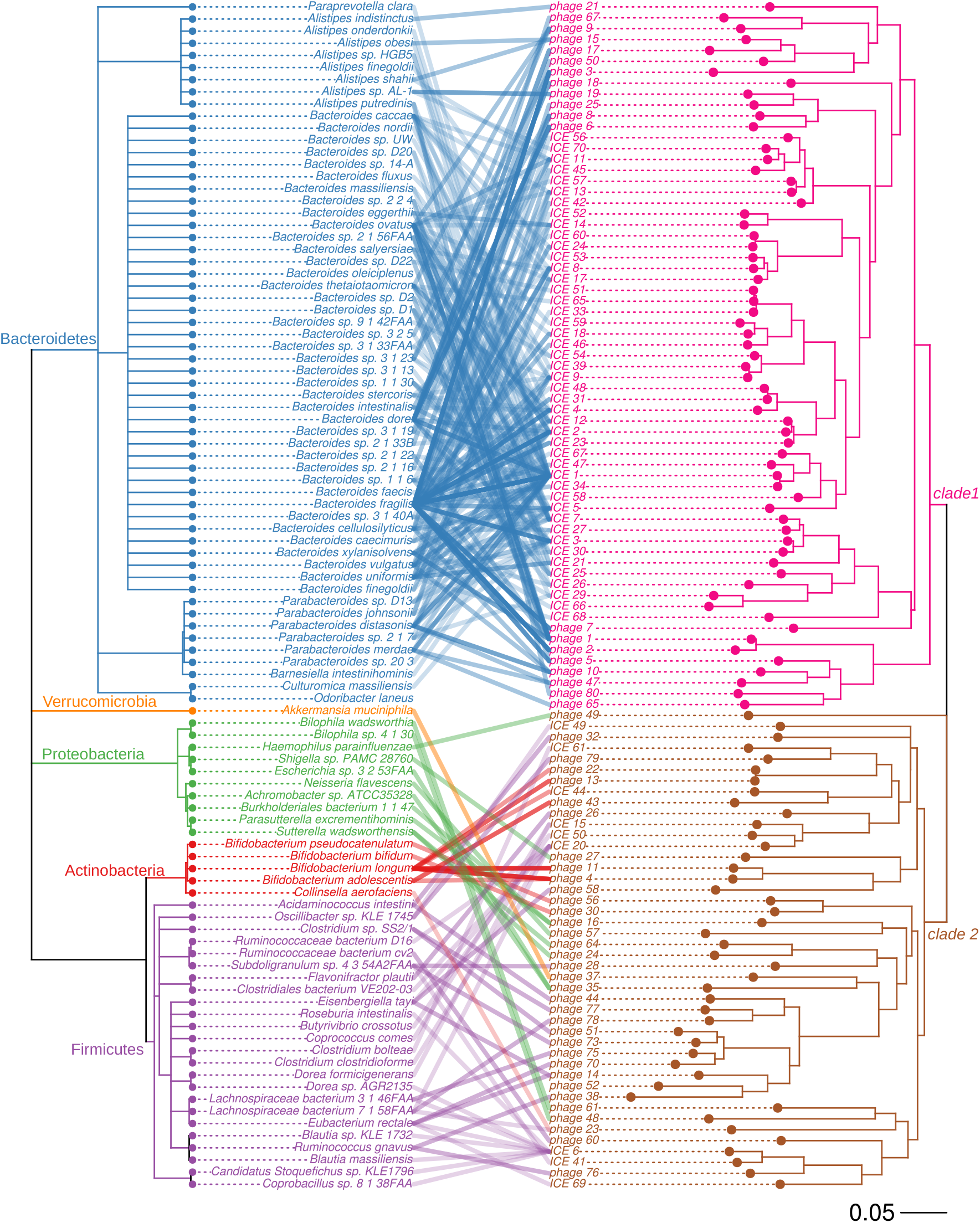
Tanglegram of host species lineages and phylogeny of the integrases in prophages and ICEs. A tanglegram of tyrosine integrases from ICEs and prophages with the species phylogeny plotted on the left and tyrosine integrase phylogeny plotted on the right. Connections are drawn between a species and the tyrosine integrase(s) found in that species and each connecting line is colored according to host bacteria phylum.

**Figure 4.**
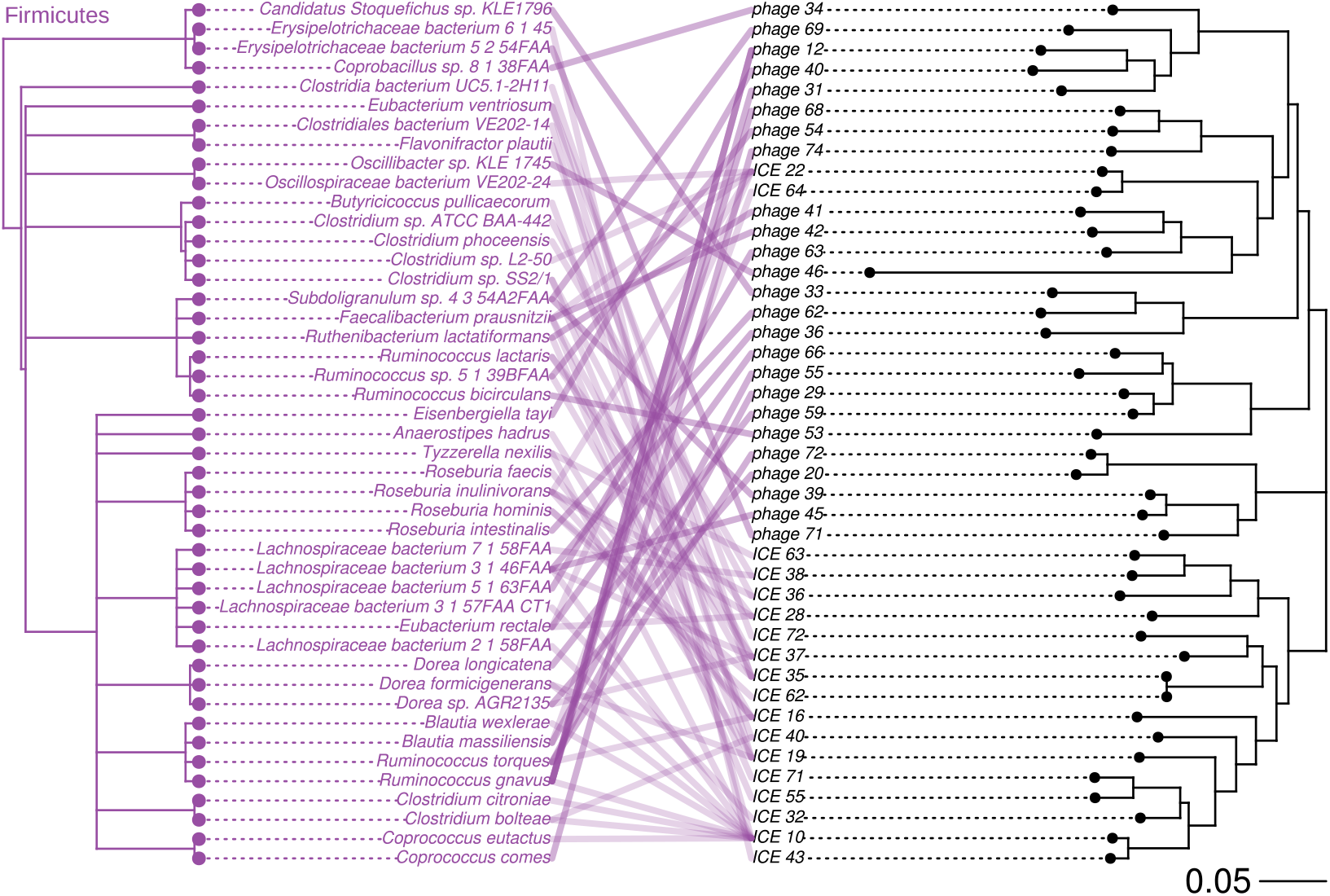
Tanglegram of host species lineages and phylogeny of serine integrases in prophages and ICEs. A tanglegram of serine integrases from ICEs and prophages with the species phylogeny plotted on the left and tyrosine integrase phylogeny plotted on the right. Connections are drawn between a species and the serine integrase(s) found in that species.

The tyrosine integrases can be divided into two clades: the first is associated with the phylum Bacteroidetes, the second clade is associated with the phyla Proteobacteria, Actinobacteria, Firmicutes and Verrucomicrobia (Figure 3). Tyrosine integrases from Bacteroidetes show no evidence of close inter-phyla transfer but ancient transfers of tyrosine integrases between the phyla Proteobacteria, Actinobacteria, and Firmicutes likely occurred several times during evolution. Serine integrases from ICEs and prophages were only found in the phylum Firmicutes. Therefore, we found no evidence of inter-phyla transfer for ICEs and prophages with serine integrases suggesting a phylum-level restriction in host range.

We also examined whether integrases derived from ICEs and prophages segregated into clades based on element type. Previous studies on the phylogenetic relationships of integrases from ICEs and prophages did not find strong evidence of intermingling between ICE and prophage integrases^56,57^. In our phylogeny of the tyrosine integrases, ICE and prophage integrases are extensively intermingled, suggesting that ICEs and prophages have exchanged integrases multiple times over the course of evolution. Moreover, in our phylogeny of serine integrases, ICE22 and ICE64 appear in a branch containing mostly prophages, suggesting that the integrase may have originated from a prophage integrase.

Unlike prophages and ICEs, 8 of 67 clusters of IMEs use serine integrases to transpose in Bacteroidetes. This implies that although integration via serine integrases occurs in Bacteroidetes, it occurs much less frequently than integration via tyrosine integrases. For transposons, 17 out of 19 transposase families were found in species from different phyla, indicating an extensive history of ancient horizontal transfer. Based on the tanglegram of group II intron proteins, no phylum corresponds to a single clade of group II introns, indicating cross-phyla horizontal transfers during the evolution of group II introns in the gut microbiome (Supplementary Figure 2).

In summary, although ancient cross-phyla horizontal transfers did occur during the evolution of MGEs, we did not observe recent cross-phyla horizontal transfer of MGEs. Therefore, the gene pools that are shared within the gut microbiome are likely limited to the phyla-level.

### Modular evolution of gut MGEs

Genes in MGEs are typically organized in functionally related modules which can be readily exchanged between MGEs. Type of modules found in MGEs include: conjugation, integration, regulation, and adaptation. Deletion, acquisition, and exchanges of these modules can lead to immobilization, adaptation, and shifts in insertion specificity and host ranges of MGEs^13^. Here, we detail examples of each of these types of events.

Many unclassified genetic islands are likely remnants of ICEs or prophages due to the presence of only a subset of genes necessary for autonomous transfer. In many cases, the integrase have been lost while other genes for conjugation or capsid formation are maintained. One example is GI73, which appears to have formed when a CTnDOT-like element lost its conjugation and mobilization modules to a large deletion (Figure 5A). We also observed many examples of the acquisition of new modules by insertions. CTnDOT-like elements have obtained adaptive modules via insertions of a group II intron together with the antibiotic resistance gene ErmF, an IME containing multiple antibiotic resistance genes including: ANT6, tetX, and ErmF^54,58^, and other unidentified insertions containing many antibiotic resistance genes (Figure 2A; Supplementary Data 3; Supplementary Table 5). Other examples are GI90, where ICE7 (CTnBST) inserted into a CTnDOT-like element (Figure 5A), and GI46, a genomic island formed when two types of ICEs (ICE43 and ICE56) inserted in tandem (Figure 5B). We observed that the exchange of recombination modules is common. Integrases have frequently been exchanged between ICEs and prophages during the evolution of MGEs (Figure 3 and Figure 4). Exchanges also occur in the same class of MGE. For example, we observed that two clusters of ICEs, ICE15 and ICE16, share nearly identical sequences and the same typeFA conjugation module, but have different integrases: ICE15 has a tyrosine integrase while ICE16 has a serine integrase (Figure 5C). Overall, the modular nature of MGEs enables the formation of new mosaic elements, leading to the diversification of MGEs, and increasing the dynamics of the gene pools in the gut microbiome.

**Figure 5.**
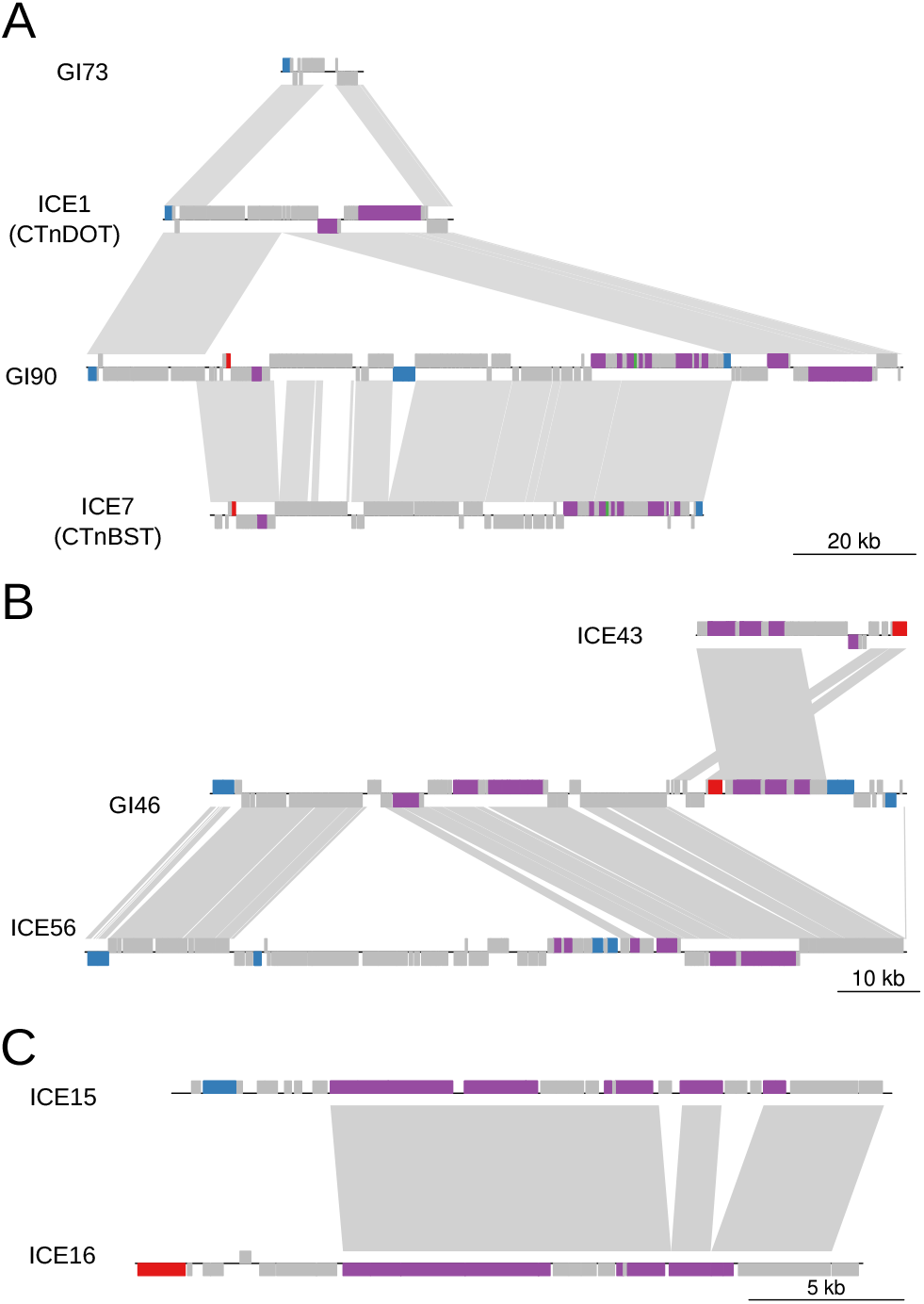
Modular evolution of MGEs. Examples of deletion, acquisition, and exchange of gene modules between MGEs. Orthologous genes between elements are shown with grey connections and are plotted with genoPlotR. Tyrosine integrases are colored blue, serine integrases are colored red, and genes involved in conjugation are colored purple. (A) GI73 was likely formed via a deletion of a CTnDOT-like ICE. GI90 was formed from an insertion of the ICE CTnBST into a CTnDOT-like ICE to form a large, composite GI that transfers as a unit. (B) An example of the tandem insertion of two ICEs to form a larger GI that moves as a unit. (C) An example of recombination module exchanges between ICE15 and ICE16.

## Discussion

In this study, we systematically characterized MGEs from the gut microbiome using a novel method to identify the mobilizable unit of active MGEs. We dramatically expanded the number of annotated MGEs from gut microbial species by identifying 5600 putative MGEs. The MGEs we identified allows for the understanding of several fundamental questions about the role of MGEs and their importance to the evolution of species of the gut microbiome.

### Implications for future gut metagenomic analysis

The database of MGEs we have curated will be a valuable resource for future studies on the gut microbiome, especially with the increasing importance of taxonomic profiling, strain-level variation detection, and pangenome analyses. Many metagenomic workflows for taxonomic profiling use marker genes or k-mers “unique” to a specific species, where uniqueness is constrained by the available reference genomes^59–61^. These marker gene should exclude MGEs, as the potential horizontal transfer of these elements invalidates their “unique” species-specific associations. Strain-level variation analyses that based on single nucleotide polymorphisms (SNPs) or copy number variation should also exclude SNPs from MGEs^62–65^. In pangenome analysis, it is beneficial to distinguish the accessory genes unique to an individual species and the mobilome shared among multiple species. To address the problems posed by MGEs to metagenomic workflows, an approach common in eukaryotic genomics, repeat masking, can be applied^66,67^. The database of curated MGEs identified in this study can be used to mask gut microbiome reference genomes before metagenomic workflows such as species-level classification, strain-level detection, and pangenome analyses are performed.

### Host ranges of MGEs and the spread of antibiotic resistance genes

In the United States alone, more than 23,000 people die each year from antibiotic-resistant infections^68^. Tracking antibiotic resistance is one of the key actions to fight the spread of antibiotic resistance. The human digestive tract is a major reservoir of antibiotic resistance genes and likely serves as a hub for the horizontal transfer of antibiotic resistance genes from commensals to pathogens^4,5,53^. MGEs play a significant role in the spread of antibiotic resistance genes, and we found that many MGEs in the gut microbiome contain antibiotic resistance genes. This study helps to define the host range of MGEs in the gut microbiome. Our results suggest that HGT occurs mostly within a phylum, and inter-phyla HGT is rare. These results underscore the risk posed by transfer of antibiotic resistance genes like the vancomycin-resistance conferring gene VanB between commensal Firmicutes and pathogenic Firmicutes, such as *Enterococcus faecalis*^69^. Overall, our study advances the understanding of the host range of MGEs which is of critical importance to understand gene flow networks in the gut.

This study underestimates the extent of host range because only MGEs in sequenced genomes were detected. As more bacterial genomes are sequenced, the extent of host range of MGEs will be refined. The scope of our research is chromosomal MGEs. Thus, plasmids or prophages existing as an extrachromosomal plasmid were not characterized in this study. Future studies using a combination of molecular and computational approaches are beneficial to further understand the rate and extent of horizontal gene transfer by MGEs.

### Niche-adaptive genes in the communal gene pool

The mammalian gut is a unique ecological niche vastly different from other environments due to the presence of IgA, antimicrobial peptides, bile acids, as well as specific polysaccharides available for utilization in the intestinal mucus. The microbes that inhabit the gut must develop mechanisms to cope with these challenges. We observed that MGEs transfer genes to help address the unique challenges of colonizing the human gut. MGEs influence the spread of gut adaptive genes in three ways. First, the spread of MGEs drives the expansion and diversification of protein families such as those involved in polysaccharide utilization, capsular polysaccharide biosynthesis, and sensing and responding to the environment^9^. Second, MGEs transfer successful innovations for colonizing the gut among distantly-related species from the same niche, such as bile salt hydrolases. Third, MGEs allow for the amplification and transfer of genes that are adaptive only under specific conditions, such as antibiotic resistance genes, and sporulation-related genes.

Cargo genes transferred by MGEs can have wide-ranging effects on the biology of the gut microbiome. They potentially involved in bacterial symbioses, sensing and responding to environmental stimuli, and metabolic versatility. The enriched classes of cargo genes we identified in this study are attractive targets for future studies to understand the underlying biology of the gut microbiome.

### Opportunities to use MGEs to engineer gut microbes

Tools for genome editing only exist for a very limited number of species of the gut microbiome despite the exceptional basic and translational opportunities afforded by engineering gut species. Many of the tools for editing the genomes of species were originally derived from MGEs. For instance, the NBU system used to modify some *Bacteroides* species was originally derived from an IME^70^, and the TargeTron system was originally derived from a group II intron^71^. The novel examples of MGEs identified in this study could be used to edit genomes from the gut microbiome, especially in currently intractable species such as *Faecalibacterium prausnitzii*. Unlike phages, whose cargo genes are limited by the capsid size, many novel ICEs and IMEs carry hundreds of genes that can confer selective advantages for the host, and are excellent candidate vectors for large genetic loci. Overall, the MGEs identified in this study could have translational applications for genome editing of species from the gut microbiome.

## Methods

### Detection of putative MGEs

80 Samples from the Human Microbiome Project (HMP)^21^ and 66,232 bacterial genomes were downloaded from the NCBI (2016/09/14). We used Mash^72^ to calculate the minhash distance between each genome and all metagenomic samples with the default sketch size of s = 1000 and *k* = 21. If the matching-hashes shared between a genome and the 80 metagenomic samples are less than 2, the genome is unlikely have enough alignments from these samples and was removed. This steps help us quickly remove genomes that likely do not exist or exist in low abundance in gut microbiome. 9,846 genomes remained after this filtering step. Metagenomic reads from HMP samples were aligned to each of the 9,846 genome separately with bwa (version 0.7.5a-r405)^73^. To find genomic regions that differ in terms of insertions/deletions between strains in the individual samples and the reference genomes, we used split reads and information from pair-end reads from the alignment (Figure 1A). First, we identified putative deletion junctions using split reads, which we defined as reads that align to two distinct portions of a genome. Split reads were initially identified as those reads having multiple hits in the SAM output from bwa. If a split read alignment starts at one genomic location in the reference and then “jumps” to aligning to a distant site downstream in the same strand, it may indicate a potential deletion in the strain of bacteria from the metagenomic sample compared to the reference genome. For each putative deletion junctions, we confirmed the presence of the junction by determining if paired-end reads flanked the junction. We considered a deletion junction to be valid if the reads pairs flanking the junction were aligned in the correct orientation, and the distance between the pairs minus the junction size is within the range of +/- 2 times the standard deviation of the mean insertion size (202.4 +/- 2X71.5 for our data set). Regions with more than four split reads and more than four read pairs supporting the deletion were considered as putative MGEs. We chose the MGEs ranging in size between 1kbp and 150kbps to reduce the number of spurious results. In total, we identified MGEs in 703 genomes. The code used to implement the SRID method, genome assembly accession numbers and HMP SRA accession numbers used in this study are available from github (https://github.com/XiaofangJ/SRID) and the Supplementary Data 4.

### MGE signature detection

Genes from the 703 genomes identified before were predicted with Prodigal (version 2.6.3)^74^. Protein sequences were functionally annotated with interproscan (version 5.19-58.0) using the default settings^75^. Then, we used the interprosan annotations to identify serine and tyrosine integrases as well as group II intron proteins from all genomes. prophage-related genes were identified by searching for genes with Pfams signatures identified in phage_finder^76^. Serine integrases were identified as genes annotated with the Pfam identifiers: PF00239 (Resolvase: resolvase, N terminal domain), PF07508 (Recombinase: recombinase), and PF13408 (Zn_ribbon_recom: Recombinase zinc beta ribbon domain). Tyrosine integrases were identified as genes annotated with the identifiers: PF00589 (Phage_integrase: site-specific recombinase, prophage integrase family), PF02899 (Phage_integr_N: prophage integrase, N-terminal SAM-like domain), PF09003 (Phage_integ_N: bacteriophage lambda integrase, N-terminal domain), TIGR02225 (recomb_XerD: tyrosine recombinase XerD), TIGR02224 (recomb_XerC: tyrosine recombinase XerC), and PF13102 (Phage_int_SAM_5: prophage integrase SAM-like domain). Group II intron proteins were identified as genes annotated with the identifier: TIGR04416 (group_II_RT_mat: group II intron reverse transcriptase maturase). To identify genes in involved in mobilization and conjugation of MGEs, we used ConjScan via a Galaxy web server (https://galaxy.pasteur.fr/)^77^. We identified transposases using blastp against the IS database with an e-value 1-e3^17^. The best hit for each protein was used to annotate the family of transposases.

### Classification of MGEs

Putative MGEs were annotated as an ICE if they contained complete conjugation and relaxase modules and an integrase or DDE-transposase at the boundary of the element. Putative MGEs were annotated as prophages if there is an integrase or DDE-transposase at the boundary of the element and more than five genes were annotated with prophage-related Pfams. Putative MGEs were annotates as IMEs if they contained an integrase or DDE-transposase and relaxase did not contain genes involved in conjugation. Putative MGEs were annotated as transposons if they contained transposase and were not previously annotated as an IME. We limited the size of IMEs to 30kb and transposons to 10kb to decrease the number of false positives. Putative MGEs were annotated as group II introns if the element was less than 10kb, contained a protein with the TIGR04416 signature, and did not contain a gene annotated as transposase. The remaining putative MGEs were then divided into two groups based on their sizes: unclassed genomic islands (>10kb), and islets (<10kb). To eliminate spurious MGEs, we only report genomic islands that contain an integrase or DDE transposase, or those that are related to prophage/ICEs, and islets that exist in more than two species. After classification and verification, we identified 5600 MGEs in 542 genomes(Supplementary Data 1;Supplementary Data 2).

### Clustering each class of MGEs

Pairwise alignment of elements from the same class of MGEs was performed with nucmer (version 3.1)^78^. Elements with more than 50 percent of the sequence aligned to each other are grouped in the same cluster. For ICEs, we additionally require that elements in the same cluster should have the same types of integrase, relaxase and conjugation modules. For IMEs, we required that each cluster has the same the types of integrases and relaxases for all elements. For transposons, the same cluster should have the same type and number of IS genes. If a transposon is a “nested” or composite transposon, the family names of all IS contained within were used to annotate the transposon.

### Construction of phylogenetic trees

To build phylogenetic trees of ICE and prophage integrases, we selected a representative integrase sequence for each cluster. For group II introns, we selected a representative group II intron reverse transcriptase/maturase from each cluster.The representative protein is a single protein chosen that has the greatest amino acid identity, on average, to its homolog sequences of the same cluster. We performed alignment of each group of sequences with mafft(v7.123b)^79^ (parameter “--maxiterate 1000”). We used trimal (version 1.4.rev15)^80^ to remove region with gaps representing more than 20% of the total alignments (parameter “-gt 0.8”). RAxML(version 8.2.10)^81^ was used to build the phylogenetic trees from the alignments using the LG substitution matrix and a gamma model of rate heterogeneity (parameter “-m PROTGAMMALGF”). Phylogenetic trees were plotted with the R package phytools^82^.

### Functional enrichment analysis of cargo genes

Cargo genes are identified by excluding genes involved in transposition and transfer from all genes on MGEs.

To understand the function of cargo genes, we performed enrichment analysis based on gene ontology (GO), antibiotic resistance (Resfam), and protein families (Pfam). The enricments were performed with all genes present in the genomes as background reference. We used hmmer^83^ to search Resfam^28^ database to annotate antibiotic resistant gene. The “--cut_ga” parameters were used to set the threshold. The best hits to each gene from the Resfam database were used to annotate antibiotic resistant genes. GO terms and Pfam signature of the same genes sets were extracted from interproscan result. R package GOStat^84^ was used for GO enrichment analysis for GO and Pfam. The R package clusterProfiler^85^ was used for the enrichment analysis of cargo genes based on Resfam and Pfam signatures. P-value of 0.05 were used as cutoff for all enrichment analysis.

## Acknowledgements

Acknowledgements

This work is supported by the Center for Microbiome Informatics and Therapeutics at MIT. A.B.H. is a Merck Fellow of the Helen Hay Whitney Foundation.

## Competing financial interests

Eric Alm is a co-founder and shareholder of Finch Therapeutics, a company that specializes in microbiome-targeted therapeutics. Other authors declare that they have no competing interests.

## Authors' contributions

Data generation, analysis, and presentation: XJ and ABH; Writing of the manuscript: XJ, ABH; Initiated the study, provided resources, tools and critical review of manuscript: RJX, EA. All authors read and approved the final manuscript.

## Supplementary information

**Supplementary Figure 1:** Classification of gut microbiome MGEs at the phylum level

**Supplementary Figure 2:** Tanglegram of host species lineages and phylogeny of group II intron proteins

**Supplementary Table 1:** Antibiotic resistant genes identified in MGEs classes

### Supplementary Tables and Data

**Supplementary Table 2:** Annotation and classification of MGEs (xlsx)

**Supplementary Table 3:** Niche adaptive cargo genes (xlsx)

**Supplementary Table 4:** Enrichment analysis of cargo genes (xlsx)

**Supplementary Table 5:** Annotation of regions with ARG insertions in CTnDOT-like elements (xlsx)

**Supplementary Data 1:** MGE sequences (fasta)

**Supplementary Data 2:** Annotation of genes in MGEs (gff3)

**Supplementary Data 3:** Sequences of CTnDOT-like elements with ARG insertions (fasta)

**Supplementary Data 4:** Scripts to implement the SRID method and the genome assembly accession numbers and HMP SRA accession numbers used in this study.(zipped txt)

